# Inversion and computational maturation of drug response using human stem cell derived cardiomyocytes in microphysiological systems

**DOI:** 10.1101/366617

**Authors:** Aslak Tveito, Karoline Horgmo Jæger, Nathaniel Huebsch, Berenice Charrez, Andrew G. Edwards, Samuel Wall, Kevin E. Healy

## Abstract

While cardiomyocytes differentiated from human induced pluripotent stems cells (hiPSCs) hold great promise for drug screening, the electrophysiological properties of these cells can be variable and immature, producing results that are significantly different from their human adult counterparts. Here, we describe a computational framework to address this limitation, and show how *in silico* methods, applied to measurements on immature cardiomyocytes, can be used to both identify drug action and to predict its effect in mature cells. Our synthetic and experimental results indicate that optically obtained waveforms of voltage and calcium from microphysiological systems can be inverted into information on drug ion channel blockage, and then, through assuming functional invariance of proteins during maturation, this data can be used to predict drug induced changes in mature ventricular cells. Together, this pipeline of measurements and computational analysis could significantly improve the ability of hiPSC derived cardiomycocytes to predict dangerous drug side effects.

## 1 Introduction

The discovery of human induced pluripotent stem cells (hiPSCs) has started a new era in biological science and medicine. These reprogrammed somatic cells can be differentiated into a wide variety of cell lineages, and allow *in vitro* examination of cellular properties at the level of the human individual. In particular, this technology has large implications in drug development, moving us away from well studied but often unrepresentative animal models towards direct testing of compounds in specific human phenotypes and genotypes. This new access offers the potential for creating more cost effective, better, safer drug treatments; both from the ability to target precision, patient specific approaches, and to reveal possible side effects of drugs in the broader human population. However, despite its promise, the technology needed to fully utilize hiPSCs for drug testing is still under development and currently faces many difficulties limiting practical applicability.

In particular, the problem of *maturation* is a major challenge to the successful use of hiPSCs in drug discovery and development. Although hiPSCs can be used to create specialized human cells and tissues, these rapidly grown cells and tissues may have significant proteomic and structural differences to, and are often more fetal-like than, their adult *in vivo* counterparts. This is especially true in hiPSC derived cardiomyocytes (hiPSC-CMs), where the adult cells they are intended to represent have undergone decades of growth and development under cyclical physiological loading and stimulation. However, despite this limitation, hiPSC-CMs have already been successfully used to assess unwanted side effects of drugs (see e.g., [1, 2]), and new technologies such as microphysiological systems (MPS) [3], are emerging to improve maturation and better capture drug effects. Still, the overall applicability of hiPSC-CMs to find unwanted side effects of drugs for adult cardiomyocytes remains limited by the fact that only relatively immature cells are available for analysis (see e.g., [4, 5, 6, 7]). And, as pointed out in numerous papers (e.g., [8, 9, 10, 11, 12]), the electrophysiological characteristics of hiPSC-CMs and adult cardiomyocytes differ significantly and, for determining potential dangerous drug side-effects, these differences may lead to both false positives and false negatives (see e.g., [13, 3]).

Meanwhile, *In silico* methods for investigating the properties of the action potential (AP) of excitable cells is a well-developed field (see e.g. [15, 16, 17]) and includes models of human cardiomyocytes (see e.g., [18, 19, 20, 21]), and models where the effect of drugs are taken into account (see e.g., [22, 23, 24]). Also, mathematical models of the action potential of hiPSC-CMs have been developed (see e.g., [9, 25]) based on measurements reported in [8, 26, 27, 28]. This field has progressed to the point where computational models are now an active part of cardiotoxicity research [29], and are being integrated into guidelines for comprehensive drug arrhythmia analysis.

In this work, we discuss how computational models of immature (IM) and mature (M) cardiomycytes can contribute to the improvement of the applicability of exploiting hiPSCs in the drug development pipeline. Despite remarkable progress in handling hiPSC-CMs under lab conditions (see, e.g [14]), the ability to create fully mature hiPSC-CMs for drug screening is likely to remain a significant challenge. In the present report, we therefore address how *in silico* computational modeling can be used to deduce properties of mature (adult) cardiomyocytes based on two real time measurements of their immature counterpart.

A key idea in our approach is that individual proteins are functionally invariant under maturation. Therefore, maturation is multiplication in the sense that, for every type of protein, the number of proteins multiply during maturation, but the function of every protein remains unaltered. In addition, the surface area of the cell and the cell volume also increase significantly during maturation, leading to large changes in current densities between the IM and M cells. The invariance of the functional properties of the IM and M versions of every protein suggests a proportionality between the associated individual currents of the IM and M cells which may explain the results obtained in [12]. We use the proportionality between the individual currents to define a maturation matrix that maps the parameterization of a model of the IM cell to a parameterization of a model of the M cell.

Our approach to estimate effects of drugs on M cells based on measurements of IM cells can be summarized as follows and is shown in Figure 1:

1. A MPS system is used to collect time averaged voltage and intracellular (cystolic) calcium waveforms, both under control conditions and in the presence of drug.
2. These voltage and calcium traces are inverted in order to define a mathematical model of the membrane and calcium dynamics of the tested IM cells. The effect of the drug is reflected in terms of changes in the maximum con-ducances of ion channels in the model.
3. The IM models are multiplied by a maturation matrix in order to obtain models for the M cells. The effect of the drug for adult cells is estimated by comparing the AP models of the M cells.

**Figure 1:**
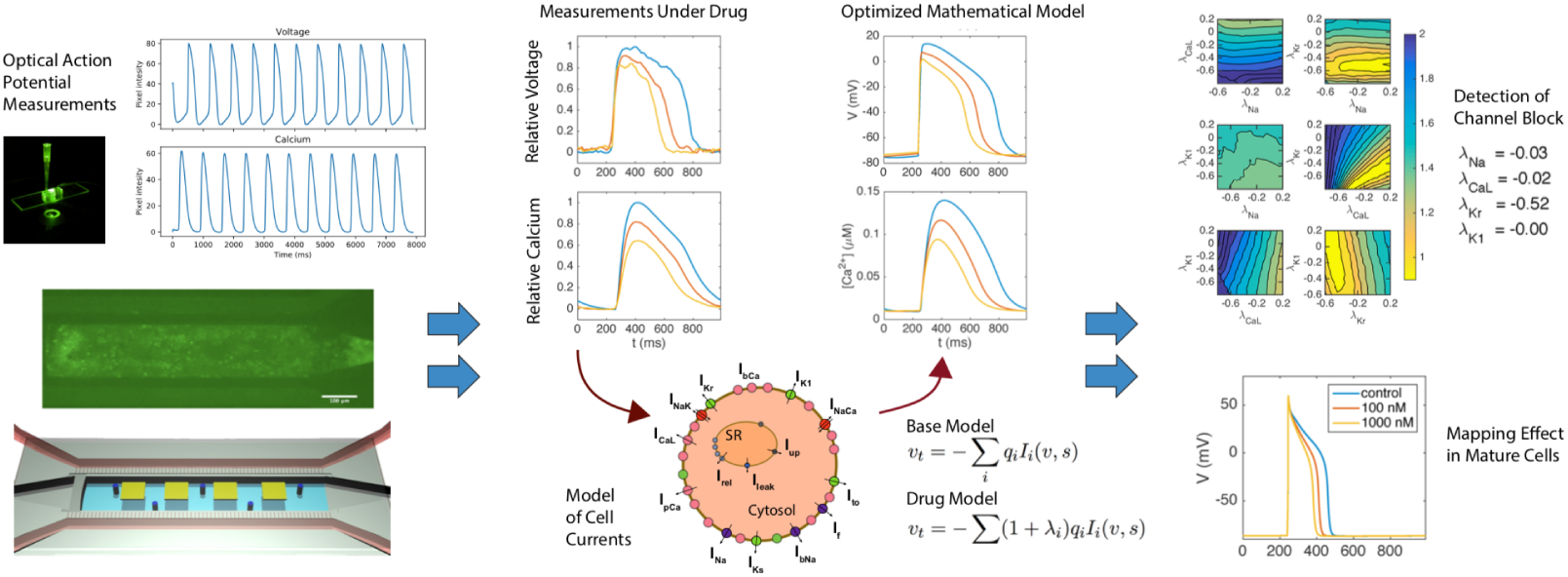
Depiction of In silico modeling and analysis of an MPS system. Optical measurements of calcium and voltage are taken at baseline and in the presence of drug. These waveforms are inverted using a mathematical model of cell dynamics, into a set of parameters that define key ion channel conductances. Changes in this parameter set give information about specific changes in conductances under drug, and this parameter set can then mapped to a model of mature cell behavior using the assumption of frunctional invariance of individual channels.

To demonstrate this process, we start by showing that a cost function, measuring the difference between data and model, is sensitive with respect to changes in the maximum conductance of major currents. Next, we show that this sensitivity is sufficient to invert simulated data and obtain a mathematical model of a drug effect. This model can be mapped from the IM case to the M case simply by multiplying a parameter vector by a diagonal maturation matrix. Finally, we apply the method of inversion to obtain an IM model based on experimental data obtained using voltage- and calcium sensitive dyes in an MPS. Again, the IM model is mapped to an M model. The effects of drugs are identified by inverting MPS data (voltage and cytosolic calcium concentration) and then mapping the resulting model from IM to M giving a mathematical model of the mature cardiomyocytes under the influence of a drug.

## 2 Results

### 2.1 Model inversion is sensitive to perturbations in major ion channel currents

The inversion of data through the minimization of a cost function requires that this cost function is sensitive to changes in model parameters. In Figure 2, we illustrate the sensitivity of three cost functions utilizing voltage, calcium, or both, to perturbations in the conductances of major cellular currents or fluxes. Here the base model (see Methods) is defined by a modified version of the Paci et al. model [9] (the details of the modification are given in the supplementary information).

**Figure 2:**
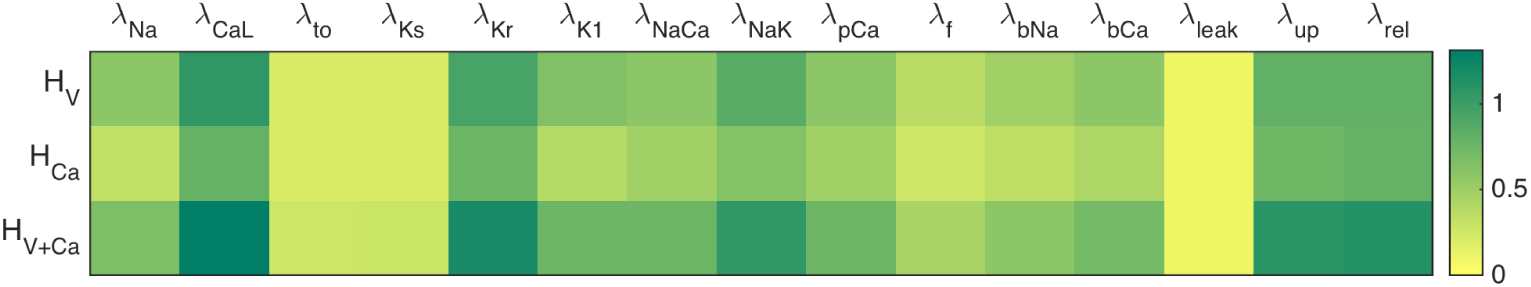
Sensitivity of maximum conductances of the immature base model assessed by the three cost functions defined in (3)–(4) with ε = 0.2. The color intensities correspond to the sum of the cost function upon reducing the maximum conductance of the given current (or flux) by ±10%.

Results indicate that cost function using voltage alone, *H_V_*, is sensitive to only some of the currents and fluxes, and in particular, it is largely insensitive to changes in *I*_to_ and *I_Ks_*. Similar trends are seen in the the calcium mismatch, *H*_Ca_, and this cost function is, in general, less sensitive than the *H_V_* version. Finally, we consider the cost function combining both the voltage and calcium data, *H_V_*_+Ca_, and observe that it is more sensitive to perturbations than both *H_V_* and *H*_Ca_ alone, although some currents are still largely invisible.

Of note the maximum upstroke velocity of the action potential is not added as a part of the *H_V_* cost function. Adding this component would likely improve sensitivity, especially for the sodium current, but our measurements (see Methods) do not at present provide sufficient accuracy of the upstroke velocity. However, the upstroke velocity of the calcium transient can be accurately estimated from the MPS measurements and is therefore a part of the cost function describing the calcium mismatch.

### 2.2 Simulated channel block identification

Although Figure 2 shows the sensitivity of the computed cost functions with respect to individual currents, we need to establish that the cost functions are adequately sensitive when multiple currents are allowed to vary. In Figure 3, we show the values of *H_V_*_+Ca_ as a function of pairwise perturbations in the maximum conductances of four major channels. The traces are theoretically computed using known effects of two chosen drugs; Verapamil which blocks *I*_CaL_ and *I*_Kr_, and Cisapride which blocks *I*_Kr_, see [29].

**Figure 3:**
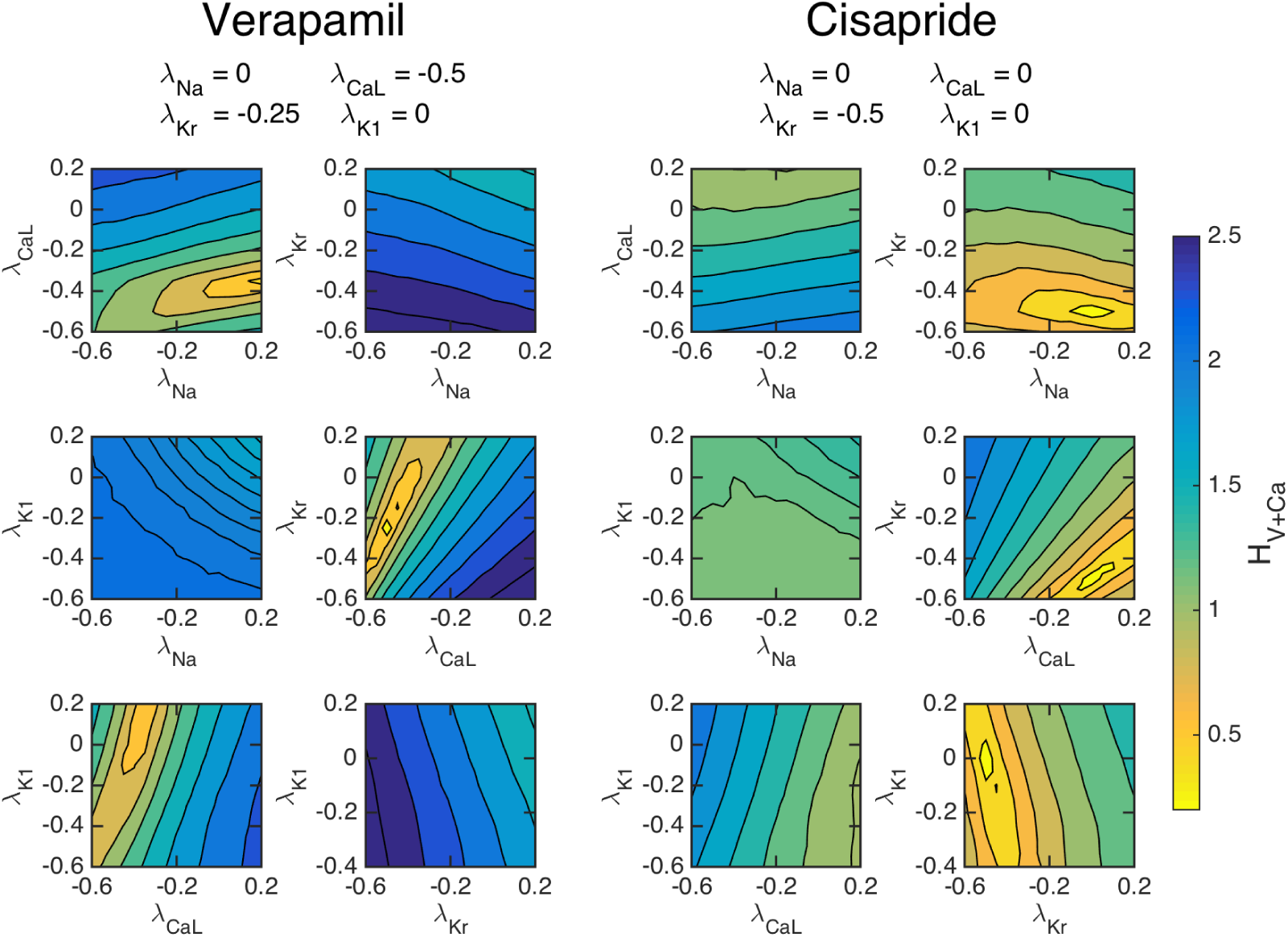
Simulated data. The cost function (4) with *ε* = 0.2 for simulated drug data, evaluated with pairwise perturbations of maximum conductances to examine if a unique minimum can be found corresponding to chosen drug effects. Left panels: The effect of Verapamil is simulated by blocking the *I*_CaL_ and *I*_Kr_ by 50% and 25%, respectively. Right panels: The effect of Cisapride is simulated by blocking the *I*_Kr_ by 50%. For both drugs, clear minimums are observed at the specified channel blockages.

Our results indicate that the cost functional using both voltage and calcium can theoretically identify the simulated channel block of the chosen drugs. The left panels show the value of *H_V_*_+Ca_ as a function of the maximum conductances when the control data are computed using the specified blocking due to the application of Verapamil. Six different configurations of pairwise blocking perturbations were tested and a minimum is clearly obtained close to the correct blocking of *I*_CaL_ and *I*_Kr_. Meanwhile, in the right panel, we show the values of *H_V_*_+Ca_ when *I*_Kr_ is blocked by 50%, simulating the effect of Cisapride. The pairwise perturbations clearly identifies that *I*_Kr_ is blocked by around 50%. These results indicate that an optimization algorithm of the cost function could find unique minima corresponding to specific channel blocks.

### 2.3 Simulated drug effect identification using the inversion procedure

Our methodology for inversion and mapping from IM to M state is first illustrated in Figure 4 using simulated data. This process is used to identify the theoretical effect of the two drugs of Verapamil and Cisapride on mature cells from waveforms that would be obtained from known channel blocking. From the left panel, we observe that the inversion algorithm is able to identify the specified effect of both Verapamil and Cisapride very accurately, reproducing chosen blocks nearly exactly. This is consistent with the results of Figure 3. The figure also shows the IM (middle panel) and M (right panel) action potentials and calcium transients. The M models are then computed using the maturation map introduced in the Methods section showing how these detected blocks would appear in mature cells.

**Figure 4:**
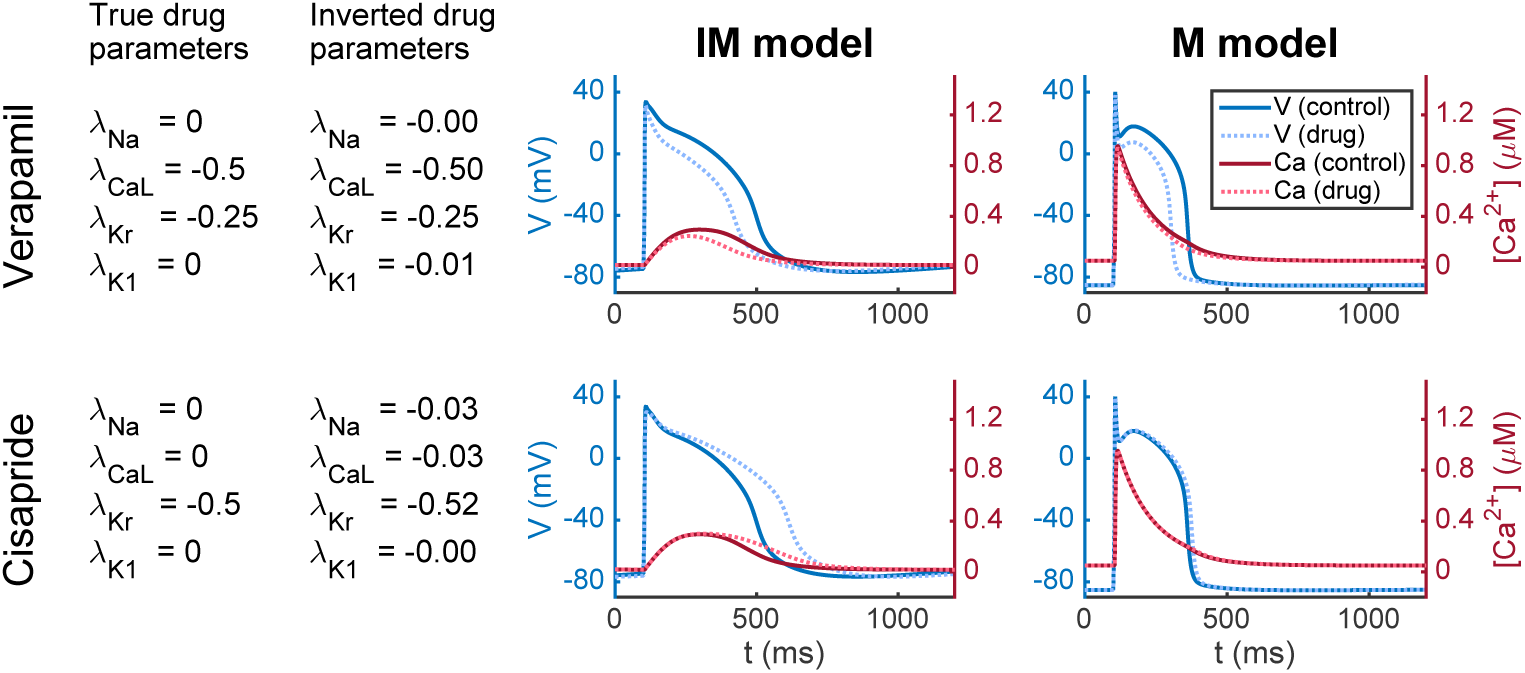
Identification of drug effects on M cells based on simulated data of IM cells. Left panel: Results of inversion by minimizing the cost function (4) with ε = 0.2. Middle panel: Action potential (blue) and calcium transient (red) before and after (dotted) the drug is applied. Right panel: Model results after application of the maturation matrix.

### 2.4 Channel block identification using a combined *in vitro / in silico system*

After demonstrating the theoretical sensitivity of inversion and drug effect prediction, we turn to the application of inverting actual cardiac MPS data. Average voltage and calcium traces (*v, Ca*) = *(v(t),Ca(t))* from an MPS [3] were inverted to yield parameterized mathematical models of the IM cells. This was done first for control data, denoted by (*v^C^*, *Ca^C^*) to yield a control model. We then show the sensitivity of the cost function *H_V_*_+Ca_ comparing this model with obtained voltage and calcium waveforms under the effect of actual doses of Verapamil and Cisparide, *(v^D^*, *Ca^D^*). In Figure 5, we present pairwise perturbations of maximum conductances and we observe again that the cost function *H_V_*_+Ca_ is sensitive to these perturbations. For Verapamil, we see that the cost function clearly indicates that *I*_CaL_ is blocked by around 50%. Furthermore, *I*_Kr_ seems to be blocked significantly, but it is not clear from the figure the extent of block. In the right panel, we also consider the effect of Cisapride. Here, the cost function indicates that *I*_Kr_ is blocked to a large extent.

**Figure 5:**
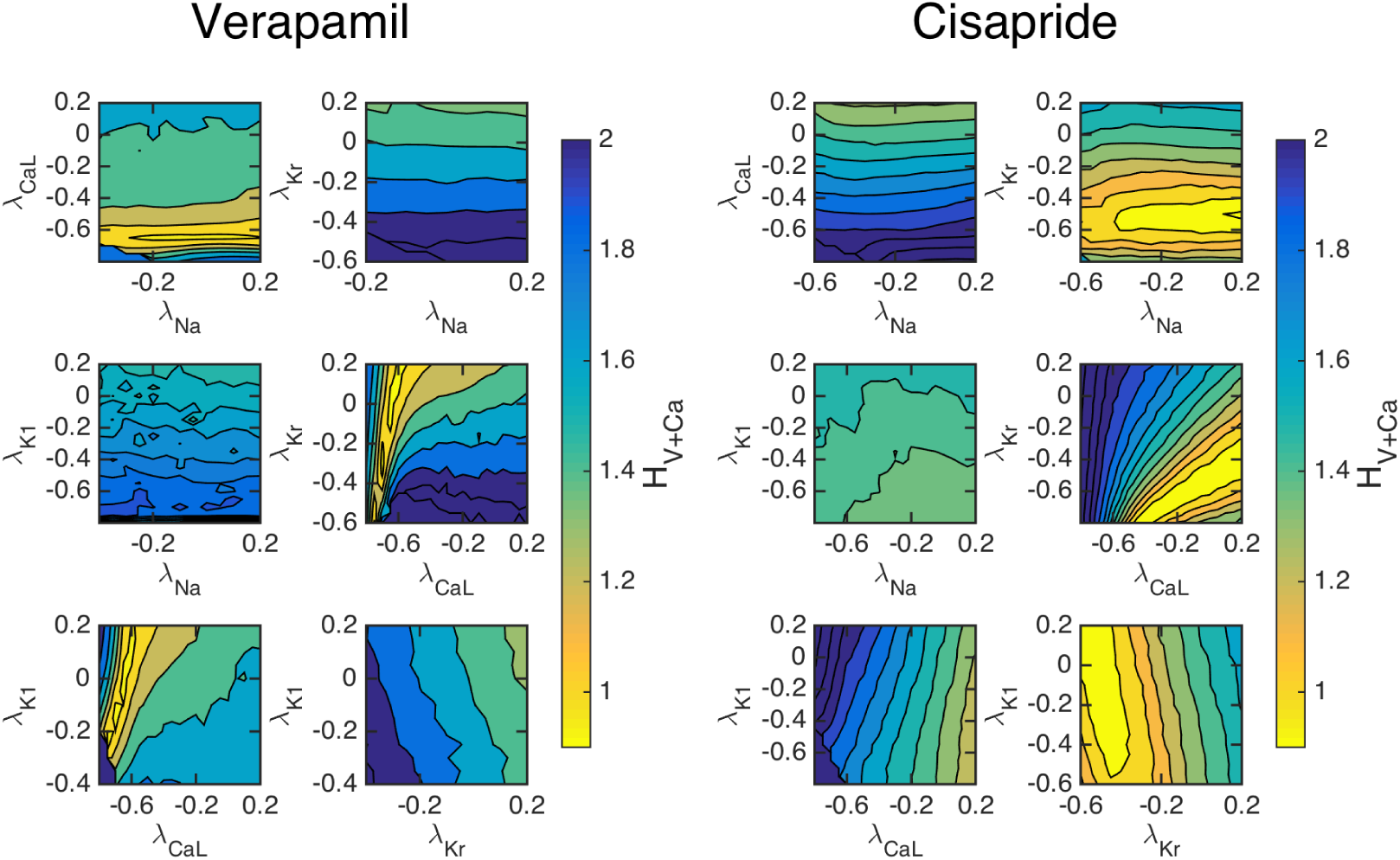
Measured data. The cost function (4) with ε = 0.2 evaluated for pairwise perturbations of maximum conductances using measured data from the MPS. Left panels: The effect of a dose of 100 nM of Verapamil is shown; it clearly blocks *I*_CaL_ and it also block *I*_Kr_. Right panels: The effect of a dose of 10 nM of Cisapride is shown; it clearly blocks *I*_Kr_. The results of the inversion is given in Figure 6.

The full inversion procedure (see the Methods section) is then applied, and it finds that *I*_CaL_ is blocked by 71% and *I*_Kr_ is blocked by 19% (see Figure 6) for Verpamil, in rough agreement with known properties of Verapamil at this dose. For Cisapride, the inversion predicts that *I*_Kr_ is blocked by 52%, and it predicts that the other currents are nearly unaffected by the drug.

**Figure 6:**
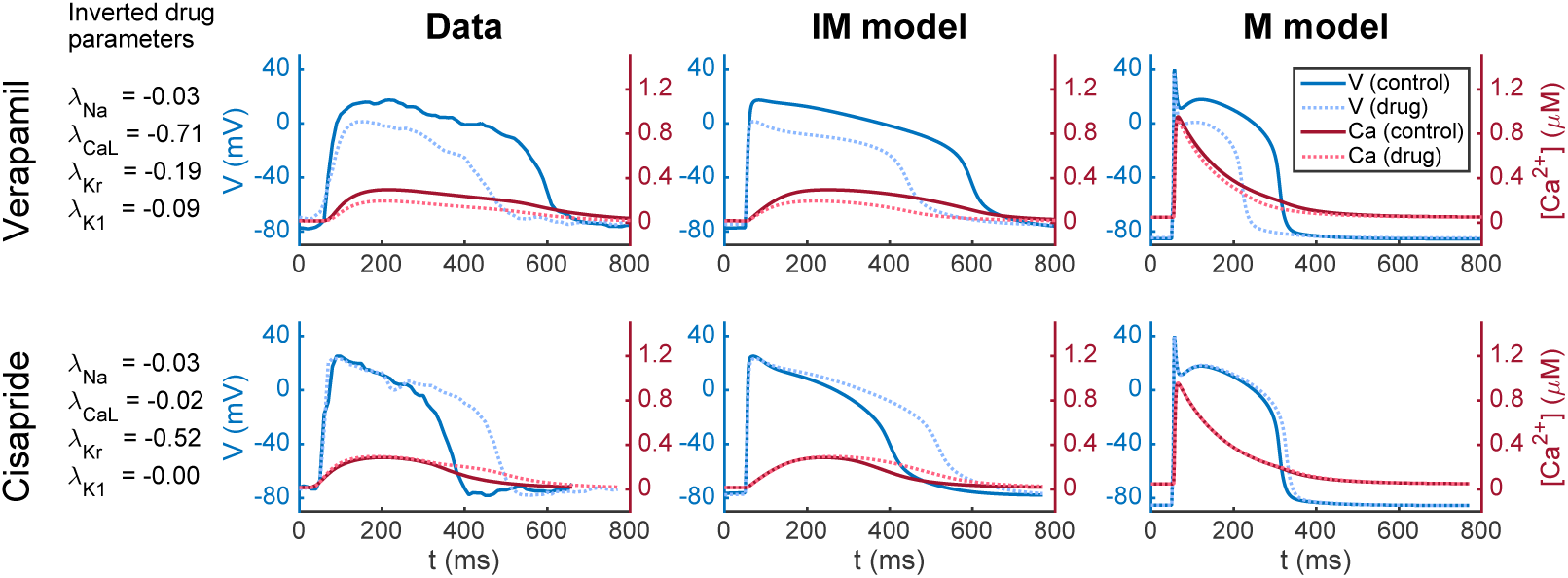
Results obtained by applying the inversion procedure to measured MPS data. First column: Results of inversion by minimizing the cost function (4) with ε = 0.2. Second column: Average voltage and calcium traces from MPS measurements. Third column: The AP model of the IM cells. Fourth column: The AP model of the M cells.

### 2.5 Mature AP change prediction using MPS data

In Panel 1 (leftmost) of Figure 6, we show the numeric results of inversion using measured data. The next three panels show action potentials and calcium transients for measured data (Panel 2), simulation of IM cells (Panel 3) and simulation of M cells (Panel 4). The simulations presented in Panel 3 are based on inversion of the MPS data giving the block values shown in Panel 1. The parameter vector (see the Methods section) representing the IM cells is multiplied by the maturation matrix in order to define the parameter vector representing the M cells. The figure illustrates how MPS measurements of IM cells can be used to estimate effect of an unknown compound for M cells.

## 3 Discussion

In this paper, we present a mathematical analysis framework to define the electro-physiologic mechanisms of drug action in mature human cardiomyocytes using only optical recordings of membrane potential and calcium in hiPSC-CMs. This novel procedure overcome a number of major existing challenges in hiPSC-CM-based screening: (1) data inversion of measured drug effects can be successfully applied to all-optical experimental data, thus allowing detailed pharmacologic characterization without the need for intracellular electrodes, (2) the mathematical approach to mapping between hiPSC-CM and adult myocyte electrophysiology is straightforward and generalizable, and (3) the MPS-based optical recordings are averaged over relatively large populations of hiPSC-CMs, thus reducing errors associated with the well-known phenotypic heterogeneity of hiPSC-CM preparations.

### 3.1 Inversion of voltage and calcium traces provide action potential models

Modern cardiac AP models have been developed more or less continually since the celebrated sinoatrial node model of Noble [30]. As a result, a range of cardiac cellular models have evolved to represent the accumulated knowledge of nearly six decades of multidisciplinary research, and the models are detailed and complex. Conventional approaches to developing these models have relied heavily upon voltage-clamp microelectrode data. These techniques provide exquisite resolution of single-channel [31, 32, 33, 34], through to whole-cell currents [35, 36, 37], and has thereby allowed the models to provide remarkably accurate reconstructions of cardiac cellular APs and calcium dynamics. However, while generalized cell models built using such data are widely used, especially to mechanistically understand how drug compounds alter electrophysiology, the experimental methods used to build them are technically challenging, have intrinsically low-throughput and cannot be used on tissue models like MPS.

In the present paper, we have developed an alternative approach that attempts to exploit the decades of information stored in modern cardiac AP models to rapidly parameterize *base models* for hiPSC-CMs. Rather than the data traditionally used to develop AP models, we used metrics that can readily be measured in a MPS, namely the optically assessed transmembrane potential and the cytosolic calcium concentration. However, these data are fundamentally different from detailed measurements of single currents traditionally used to invert measurements into biophysical models, and new methodology is needed. The approach taken in this report is based on minimization of a cost function comparing the predicted and measured waveforms, which seems to provide reasonable accuracy in analysis, but it is clear that some currents are still largely invisible even theoretically, and alternative approaches may lead to broader or more focused results.

For example, it was observed in Figure 2 that the cost function *H_V_*_+Ca_ is more sensitive than both *H_V_* and *H*_Ca_ (see (3)-(4)). This indicates that both voltage and calcium traces must be measured in order to get optimal inversion of the measurements. However, this depends on the application. For instance, if the main purpose is to study side effects on the *I*_Kr_ current, it may be sufficient to only consider voltage traces. In addition, cost functions which take into account measured extracellular potential or contractile for generated by the IM cells may also be used to better invert specific drug induced changes.

### 3.2 The maturation map

While the inversion of data from hiPSC-derived cells will be essential for understanding the electrophysiology of immature cells, understanding how such elec-trophysiolgical changes translate into mature cells could provide powerful means to screen drugs for side effects. We introduce the idea of a maturation map, which assumes that the essential difference between an immature (IM) cell and a mature (M) cell can be described by the number of proteins, the membrane area and the volume of the cell and the intracellular storage structures. Based on these assumptions, we have argued that we can map any IM parameter vector, *p_IM_*, to an associated M parameter vector, *p_M_*, simply by multiplying by a diagonal matrix *Q: p_M_ = Qp_IM_*. We have illustrated this mapping and noted that reasonable models of an IM AP are mapped over to a reasonable M AP. In addition, we have seen that measured IM data can be inverted to yield *p_IM_*, and then the maturation mapping gives the adult AP parameterized by *p_M_ = Qp_IM_*.

In the present report, we have simply addressed the mapping directly from an IM state to the M cells. However, maturation is clearly a dynamic process with rapid changes, and it may therefore be of interest to use this mapping to investigate the time dependent behaviour of the cells. Measurements of several time instances of IM cells may give insight into the developmental trajectories of IPSC-CMs and how different maturation protocols alter the electrophysiological properties of generated test cells. Such studies may be useful for both choosing maturation protocols to optimize data inversion sensitivity, and for quality control measures of batch to batch cells.

In addition, taking into account more aspects of cellular electrophysiology could refine our approach. For example, one could take into account that proteins exist in various forms; for instance, the sodium channel has nine different forms with different associated possible channelopathies. These variants may proliferate at different rates and thus potentially lead to significant changes in the properties of the M cells.

### 3.3 hiPSC data sources

While our results show the promise of this methodology, considerable current limitations exist that need to be addressed. First, variability in hiPSC-CMs remains a challenge ([38, 39]). In the preparation of the data, we have dealt with variability by discarding individual voltage and calcium traces that are significantly different from the average behaviour of the cells. This seems to give sufficient accuracy for inversion, and the effects of the drugs we have considered have shown significant cellular changes. However, even if the average results clearly respond to the doses of the drugs applied in this study, work on reducing the variability of generated hiPSC-CMs in MPSs is clearly needed for batch to batch consistency.

In addition, improvements in data acquisition from the cell systems may also improve the methodology, in particular it may increase the sensitivity of cost functions to currents that are presently less visible. For instance, the voltage waveform can not currently be imaged at the time resolution needed to obtain accurate measurements of the upstroke, due to a combination of hardware and optical light collection limitations. In the same manner, the signal to noise ratio in this waveform, due to background dye absorption, prevents adequate resolution of the plateau phase and in particular of the notch in the action potential, preventing inversion of the I_to_ current. Improvements in the methodology for collection of high resolution optical voltage data will therefore lead to substantial improvements in mapping resolution.

It should also be noted that it is possible to measure the extracellular field potential in the microphysiological systems using a multi-electrode array (MEA) system, see e.g., [40, 1]. Such data can be incorporated in our method by using the EMI model (see e.g., [41]) instead of the common AP models. In this case, the function H given by (4) would have to be extended to include the EFPs. This would be considerably more computationally demanding than the present method, but it may also increase the accuracy of the inversion.

### 3.4 Extension to specie - specie mapping

The basic idea underpinning the maturation mapping is that the proteins populating the cell membrane are the same for the IM cells and the M cells; the reason for the significant difference in AP between these cell types is the difference in densities of membrane proteins. Similarly, the proteins of the cell membranes are also quite similar from one specie to another, but again the densities of these proteins vary considerably. Therefore, the procedure for detecting side effects of drugs developed in this report *may* be generalized to be used between species. More specifically; it may be possible to measure the effect of drugs for mouse cells and deduce the effect for human cells following the steps detailed in the Method section below. This may be of significance because of the abundance of mouse data.

## 4 Methods

Our aim is to enable automatic characterization of side-effects of drugs for mature cardiomyocytes based on measurements of voltage and calcium traces of immature cells in an MPS. Here, we describe the methods applied above; we briefly how appropriate optical measurements of voltage and calcium are obtained, how a model of the AP of a mature cardiomyocyte can be obtained from a model of an immature cardiomyocyte, and how these data are inverted in order to define a mathematical model of the AP of the immature cells. Furthermore, we describe how the effects of drugs on M cardiomyocytes can be estimated using measurements of the effect on IM cardiomyocytes.

### 4.1 Measuring voltage and calcium traces using an MPS

Using previously developed techniques [3], cardiac MPS systems were loaded and matured prior to drug exposure. On the day upon which studies were performed, freshly measured drug was dissolved into DMSO (Cisapride) or media (Verapamil) and serially diluted. Each concentration of the drug to be tested was preheated for 15-20 min in a water bath at 37°C and subsequently sequentially injected in the device. At each dose, after 5 min of exposure, the drug’s response on the microtissue was recorded using a Nikon Eclipse TE300 microscope fitted with a QImaging camera. Fluorescent images were acquired at 100 frames per second using filters to capture GCaMP and BeRST-1 fluorescence as previously described. Images were obtained across the entire chip for 6-8 seconds, with cell excitation paced at 1 Hz, to capture multiple beats for processing.

Fluorescence videos were analyzed using custom Python software utilizing the open source Bio-Formats tool to produce characteristic voltage and calcium waveforms for each chip and tested drug dose. Briefly, for each analysis, the fluorescent signal for the entire visual field was averaged, excluding pixels which did not change significantly in intensity over the acquisition. The signal was then smoothed using a 3 point median filter, and 5-7 individual action potentials or calcium transients overlayed by aligning the maximum dF/dt and then averaged into a single transient.

### 4.2 Inversion of voltage and cytosolic calcium traces

In order to complete the description of the steps presented in Table 1, we need explain how the inversion used in steps 4 and 5 is performed, and the key question is how to do the inversion of the form (8). In order to explain this, we assume that we have a base model of the form

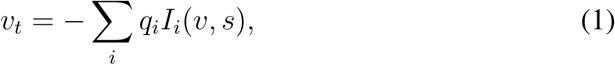

where *I_i_* represents the dynamics of the individual membrane proteins and *q_i_* represents the maximum conductance of the ion channels (or the maximum rate of an exchanger or a pump). Furthermore, *v* is the transmembrane potential and *s* represents the remaining state variables of the model. In order to adjust this model to a set of measured data given by (*v*\*, *c*\*), we seek parameters *λ_i_* such the solution of

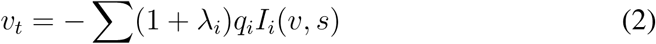

is as close as possible to the measured data, *(v*,c*).* The distance from the computed solution *(v, c) = (v(λ), c(λ))* to the measured data *(v*,c*)* is given by a cost function *H = H*(*λ*).

**Table 1:** The table shows a summary of the method for computing possible side effects of drugs for mature cells based on measurements conducted on immature hiPSC-derived cells.

|  |  |  |
| --- | --- | --- |
| 1 | Base model | $p^{M,B} = Q^B p^{IM,B}$ |
| 2 | Measure control (C) data, no drug | $(v^C, c^C)$ |
| 3 | Measure data with drug (D) applied | $(v^D, c^D)$ |
| 4 | Invert C-data | $(v^C, c^C) \xrightarrow{\text{inversion}(p^{IM,B})} p^{IM,C}$ |
| 5 | Invert D-data | $(v^D, c^D) \xrightarrow{\text{inversion}(p^{IM,C})} p^{IM,D}$ |
| 6 | Update maturation map | $Qp^{IM,C} = p^M$ |
| 7 | Parameterize M version of D cells | $p^{M,D} = Qp^{IM,D}$ |
| 8 | Compare M version of C and D cells | Simulate M cells with $p^{M,D}$ and $p^M$ |

We consider the following cost functions

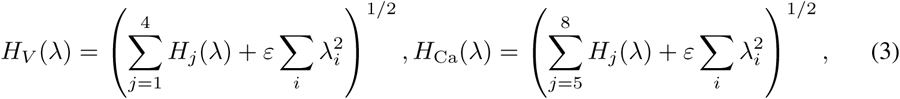

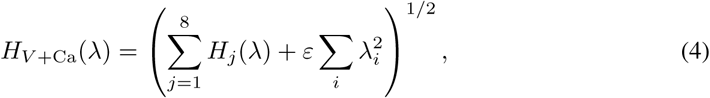

where

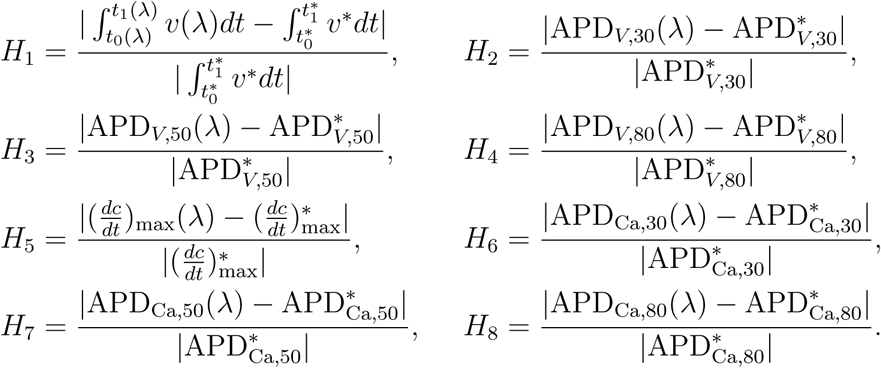

Here, the star * is used to denote observed data, either generated by simulations or gathered from the MPS. Also, 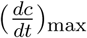 is the maximal upstroke velocity of the calcium concentration. Furthermore, APD*_V_*_,30_ is defined as the length (in ms) of the time from the value of the transmembrane potential, in the upstroke, is 30% below its maximum value (*t*_0_) until it again is repolarized to 30% of its maximum value (*t*_1_). The values APD*_V,_*_50_ and APD*_V,_*_80_ are defined similarly. Likewise, the terms APD_Ca,30_, APD_Ca,50_ and APD_Ca,80_ represent the corresponding transient durations for the calcium concentration. In *H*_1_, we compute the integral of the transmembrane potential from *t = t*_0_ to *t = t*_1_. Note that *H_V_* only depends on characteristics of the voltage trace, whereas *H*_Ca_ only depends on characteristics of the calcium trace; finally, *H_V_*_+Ca_ includes the terms of both the two former cost functions and therefore depends on the characteristics of both the voltage trace and the calcium trace.

#### 4.2.1 The minimization procedure

The inversion procedure aims to minimize the cost function measuring the difference between the target and model voltage and calcium waveforms. In every minimization, we have an existing parameter vector 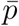, and we seek an optimal perturbation of this vector where each component is given by 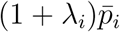. Here, *i* runs over the components of the parameter vector and *λ_i_* denotes the perturbation. The cost function introduced above is irregular and hard to minimize. Therefore, we introduce a brute force search algorithm that avoids any attempt to take the gradient into account. To start searching for suitable values of λ = {*λ_i_*}, we first set up a bounding box of allowed values of λ. This is initially set up so that each *λ_i_* is in some interval, for instance [-0.5, 0.5]. Next, we draw *N* choices of *λ* randomly from the bounding box and compute *H*(*λ*) for each of these *N* choices. We then pick out the five choices of *λ* that give the smallest values of *H*(*λ*) and set up a new bounding box of reduced size around each of these five choices of *λ*. More specifically, these bounding boxes are set up by centering the boxes around the chosen *λ* and letting the length of the interval for each *λ_i_* be reduced to 90% of the length of the previous intervals. Note that this means that the new bounding boxes are not necessarily contained in the initial bounding box, but may extend beyond the initial intervals. We do, however, set up a restriction so that no bounding box is allowed to contain values of *λ* smaller than or equal to −1. In addition, when searching for the effect of drugs, we assume that the drug is a channel blocker and therefore only consider *λ* ∈ (−1, 0].

After setting up the five new bounding boxes, we draw *N*/5 choices of *λ* randomly from each box and compute *H*(*λ*) for each of these *N* choices of *λ*. We then select the five choices of *λ* that give the smallest values of *H*(*λ*) and repeat the steps above for a given number of iterations. For the applications of the minimization method reported in the Results section, we generally use 10 iterations and *N* = 5000.

### 4.3 Maturation through multiplication

Our model of the maturation process rely on the assumption that the individual membrane proteins are functionally invariant under maturation, whereas the number of proteins, the membrane area and the cell volume change significantly (see e.g., [42, 43, 44, 11]). Also, different membrane proteins proliferate at different rates leading to large differences in the expression levels between IM and M cells. This, in turn, leads to characteristic differences between the IM and M voltage and calcium traces. The maturation process is illustrated in Figure 7.

**Figure 7:**
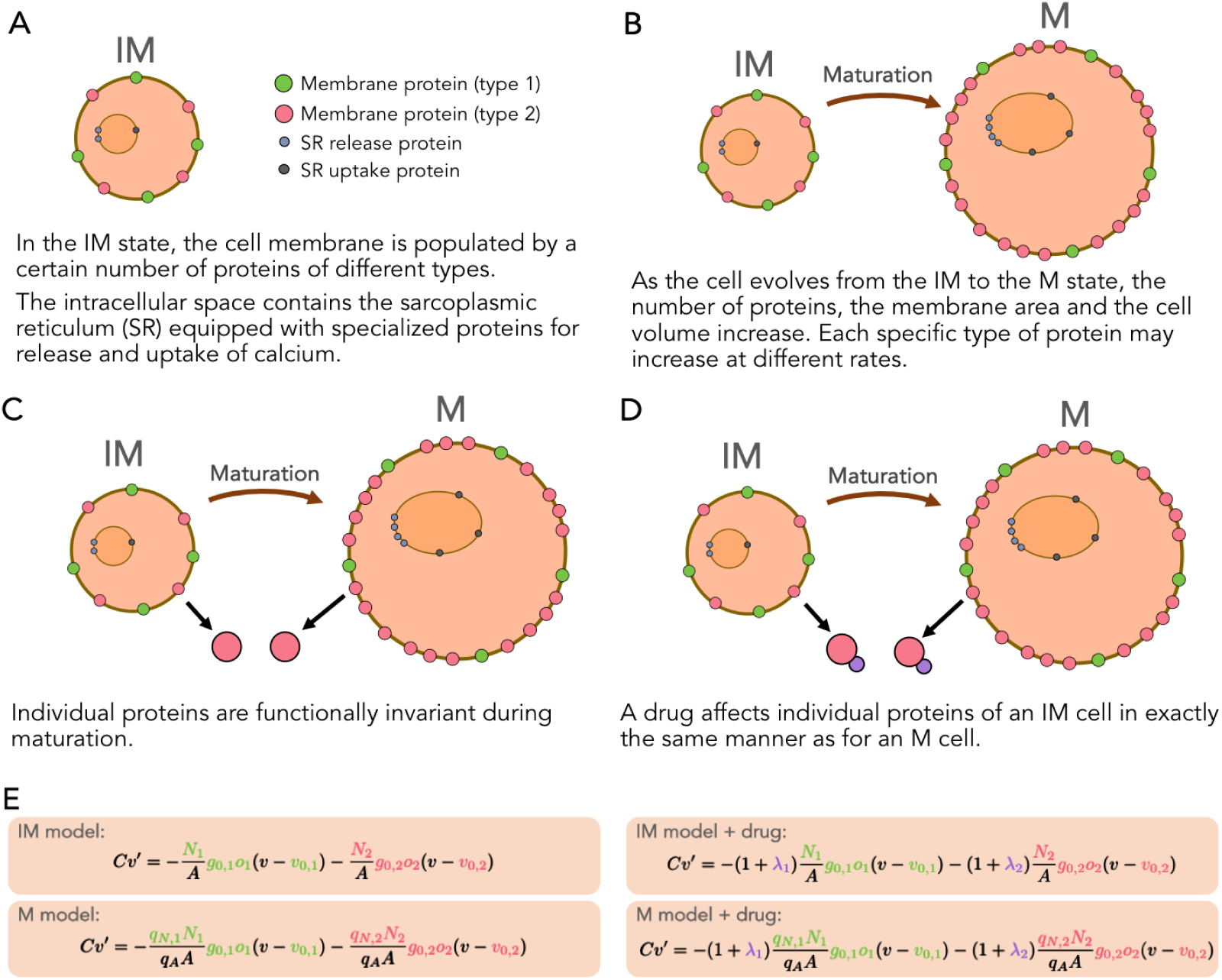
Illustration of the assumptions underlying our model of maturation. **A**: The immature cell with two types of membrane proteins, with a cytosolic space containing the sarcoplasmic reticulum with associated release and uptake proteins. **B**: Maturation is multiplication in the sense that the number of proteins increases at a protein specific rate. **C**: A specific protein in the IM cell is the same as in the M cell. **D**: A drug affects every single protein in the IM cell in exactly the same manner as for the M cell. **E**: Model of the transmembrane potential for IM and M cells, and the relation between these models; and how these models are affected when a drug is applied.

### 4.4 A drug effects a singel protein in the same manner for IM and M cells

Since we assume that exactly the same proteins are present in the IM and the M cells, it follows that the effect of a given drug on a protein in the IM case is identical to the effect on the same protein type in a M cell. This observation is essential in order to understand side effects on M cells based on measurements of the IM cells.

### 4.5 The membrane potential for IM and M cells in the presence of a single current

In order to illustrate the modeling process going from IM to M, we consider the following simplest possible case where the transmembrane potential *v* (in mV) is governed by a single current

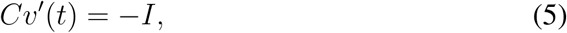

with *I = go(v − v*_0_). Here, *C* is the membrane capacitance (in *μ*F/cm^2^), *g* is the maximum conductance (in mS/cm^2^), *o* is the open probability of the channels (unitless), and *v*_0_ is the resting potential of the channel (in mV). In this formulation, the current *I* is given in units of *μ*A/cm^2^. The maximum conductance can be written on the form

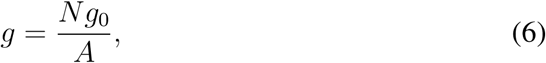

where *g*_0_ is the conductance (in mS) of a single channel, *N* is the number of channels and *A* is the membrane area of the cell (in cm^2^).

Let *N_im_* and *A_IM_* denote the number of ion channels and the surface area of the IM cell, respectively. Then there are constants *q_N_* and *q_A_* such that the number of channels in the M cell is given by *N_M_ = q_N_ N_IM_*, and the membrane area of the M cell is given by *A_M_ = q_A_A_IM_*. Therefore, the maximum conductance of the M cell can be expressed in terms of the maximum conductance of the IM cell as follows,

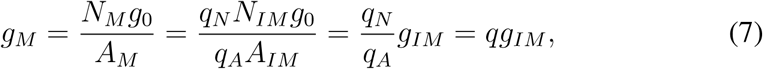

with 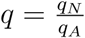.

Here, we have explained that the representation of a single current can be mapped from IM to M simply by multiplying the maximum conductance by a factor. This derivation relies heavily on the assumption that the dynamics of the single channel, represented by the open probability *o* in (6), remains the same during maturation (see Figure 7). As consequence, the Markov model (see e.g., [24]) representing the open probability of the single channel should be the same for the IM and the M version of the channel protein. Similar arguments can be presented for other membrane proteins such as exchangers and pumps. Furthermore, the intracellular Calcium machinery can be treated in exactly the same manner, leaving the IM and M models of a single protein to be distinguished only by a factor. Details of the mapping of Calcium concentration fluxes are provided in supplementary information.

The factors for the individual components of an AP model can be gathered in a parameter vector *p*, and a diagonal matrix Q can be used to store the maturation mapping from the IM parameter vector to the M parameter vector such that *p_M_* = *Qp_IM_*.

In Figure 8, we illustrate the use of the maturation mapping for well established AP models of hiPSC-CMs using the Paci et al. model [9], and of the adult human cardiomyocyte using the ten Tusscher et al. model [20]. For the Paci et al. model, we define the maturation map 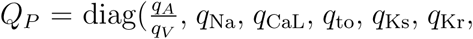 *q*_K1_, *q*_NaCa_, *q*_NaK_, *q*_pCa_, *q*_f_, *q*_bNa_, *q*_bCa_, *q*_leak_, *q*_up_, *q*_rel_) = (1.7, 0.5, 2.5, 10, 1, 0.25, 3, 0.3, 0.7, 1, 0.1, 0.32, 0.85, 200, 0.95, 35). Since *p_IM_* is given by the paper [9], we can compute *p_M_* = *Q_P_p_IM_*. Similarly, for the ten Tusscher et al. model we use 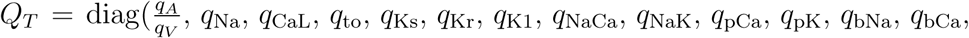 *q*_leak_, *q*_up_, *q*_rel_) = (1.7, 0.3, 2.5, 5, 10, 1.2, 50, 0.58, 0.7, 2, 0.5, 0.3, 0.85, 200, 0.6, 20), and since *p_M_* is given by the paper [20], we can compute the IM version by 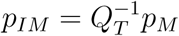.

**Figure 8:**
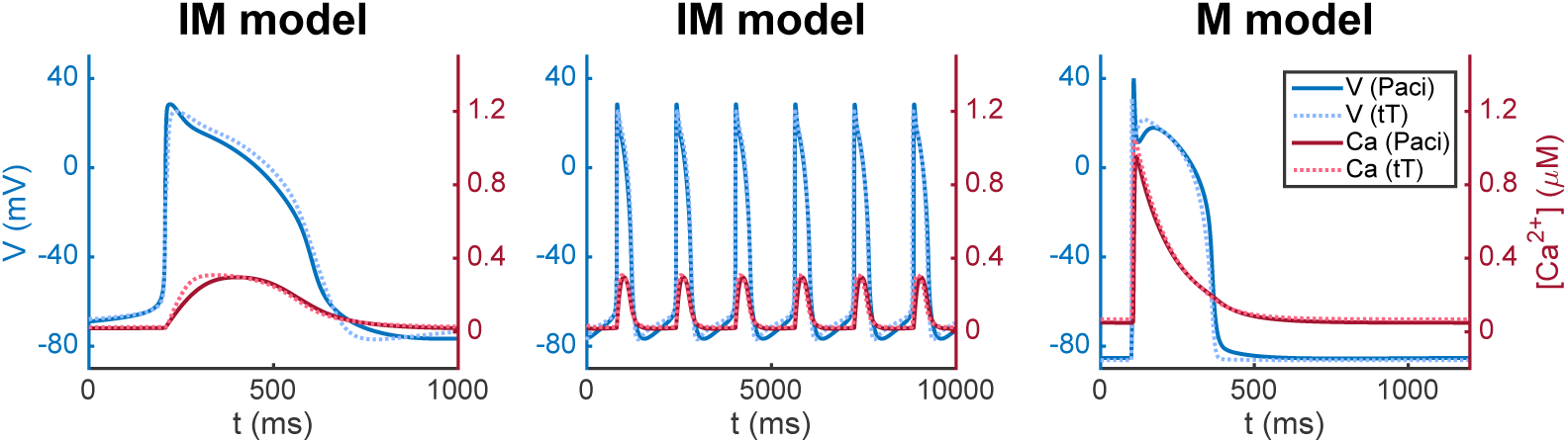
Immature and mature versions of the Paci et al. model [9] and the ten Tusscher et al. (tT) model [20]. The APs of the M cells are shorter and the upstroke velocity is faster than for the IM case; compare left and right panels. Also, the IM cells are pacemakers (middle panel), whereas the M cells reach a stable repolarized equilibrium after an AP (right panel).

We observe that these AP models have the known characteristic differences between IM and M cells; the upstroke of the IM cells are considerably slower than for the M cells, the IM cells are pacemaker cells (middle panel of Figure 8), and the M cell reaches a stable equilibrium (right panel of Figure 8).

### 4.6 Estimating side-effects drugs

The method for identifying side effects of drugs is summarized in Table 1. The method involves eight steps:

**Step 1: Base model** Assume that there exists an *AP base model,* characterized by a parameter vector *p^IM,B^*, representing a prototypical IM cell, and an associated *base maturation map Q^B^*. The associated M cells are characterized by *p^M^* = *Q^B^p^IM,B^*. The M model, parameterized by *p^M^*, provides a normal mature AP. No drug is involved in parameterizing the base model. Note also that the base model is used for numerous (independent) measurements. The base model in our computations is a modified version of the model of hiPSC-CMs suggested by Paci et al. [9]; see the supplementary information for details concerning the base model. **Step 2 and 3: MPS-measurements** For the IM cells, we measure the transmembrane potential and the cytosolic calcium concentration, stored as (*v^C^, c^C^*), and make similar measurements for the case when a drug has been applied, stored as (*v^D^, c^D^*). Here *C* is for control (no drug) and *D* is for drug. **Step 4 and 5: Inversion** Generally, the notation

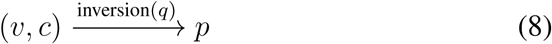

means that the data (*v, c*) are inverted to yield a model parameterized by the vector *p*, using the model parameterized by the vector *q* as a starting point for the inversion. The control data (no drug) given by (*v^C^, c^C^*) are inverted to yield the model parameterized by *p^IM,C^*, using the parameter vector *p^B^* as a starting point for the inversion. Likewise, the D-data are inverted to give the model *p^IM,D^*, where the parameter vector *p^IM,C^* is used as starting point. **Step 6: Update maturation map** The maturation map can now be updated to secure that if *Q* is applied to the IM parameter vector, *p^IM,C^*, the resulting parameter vector is the base model of the M cell parameterized by the vector *p^M^*. **Step 7: Map from IM to M** The updated maturation map *Q* is used to compute the parameterization of the M version of the drugged cells. **Step 8: Drug affected M cell** The effect of the drug on the M cells is analyzed by comparing the vectors *p^M^* and *p^M,D^*. The components of *p^M,D^* that are significantly different from its *p^M^* counterpart, has been significantly affected by the drug. The effect of the drug on the mature AP is estimated by comparing the result of simulations of the models characterized by *p^M^* and *p^M,D^*.

## 5 Acknowledgements and Disclosures

We would like to acknowledge the following funding sources: The Research Council of Norway funded INTPART Project 249885, the SUURPh program funded by the Norwegian Ministry of Education and Research, the Peder Sather Center for Advanced Study, NIH-NCATS UH3TR000487, NIH-NHLBI HL130417, and in part by California Institute for Regenerative Medicine DISC2-10090.

In addition, Professor Kevin Healy has a financial relationship with Organos and both he and the company may benefit from commercialization of the results of this research.

